# LongGeneDB: a data hub for long genes

**DOI:** 10.1101/2020.09.08.281220

**Authors:** Sohyun Moon, Yura Kim, Mariam Naghavi, Jerry Yingtao Zhao

## Abstract

The human genome contains more than 4000 genes that are longer than 100 kb. These long genes require more time and resources to make a transcript than shorter genes do. Long genes have also been linked to various human diseases. Specific mechanisms are utilized by long genes to facilitate their transcription and co-transcriptional processes. This results in unique features in their multi-omics profiles. Although these unique profiles are important to understand long genes, a database that provides an integrated view and easy access to the multi-omics profiles of long genes does not exist. We leveraged the publicly accessible multi-omics data and systematically analyzed the genomic conservation, histone modifications, chromatin organization, tissue-specific transcriptome, and single cell transcriptome of 992 protein-coding genes that are longer than 200 kb in the mouse genome. We also examined the evolution history of their gene lengths in 15 species that belong to six Classes and 11 Orders. To share the multi-omics profiles of long genes, we developed a user-friendly and easy-to-use database, LongGeneDB (https://longgenedb.org), for users to search, browse, and download these profiles. LongGeneDB will be a useful data hub for the biomedical research community to understand long genes.

## INTRODUCTION

Protein-coding genes vary in gene lengths, ranging from 0.1 kilobases (kb) to 2400 kb (1). Gene lengths affect transcription because gene transcription relies on RNA polymerase II moving across the entire gene region. Gene lengths also affect co-transcriptional processes, such as intron splicing, because a long intron results in a large RNA molecule to be removed from the mRNA precursors. Long genes require more time and resources to make a mature transcript than short genes do. Notably, long genes are linked to multiple human diseases, including amyotrophic lateral sclerosis (2,3), autism spectrum disorder (4-9), Alzheimer’s disease (10), and cancer (11,12).

Long genes use specific mechanisms to regulate their transcription and co-transcriptional processes. These specific mechanisms result in unique genomic and epigenetic features in long genes. These features include the enrichment of common fragile sites (11,12), RNA:DNA hybrids (13,14), recurrent DNA double-strand breaks (15), non-CG DNA methylation (6), intronic polyadenylation (16), and the sensitivity to topoisomerases inhibition (4). These multi-omics features are essential for researchers to understand the regulatory mechanisms of long genes and the pathophysiology of human diseases that are associated with long genes. However, these multi-omics profiles were generated by individual studies and are not easily accessible for researchers without a background in bioinformatics or computational biology. The biomedical community needs a database that can provide an easily accessible path to the ample multi-omics profiles of long genes.

The ENCODE project (17) has generated thousands of high-quality multi-omics profiles in various tissues of human and mouse. Other types of omics profiles are also available in databases such as the NCBI GEO (18) and the EBI ENA (19). These data provide a reliable and insightful resource for multi-omics data of long genes. The Ensembl database (1) contains the genomic sequences and comprehensive annotations for more than 200 species. This enables us to investigate the evolution of gene lengths of long genes in distinct species. Based on these data, we characterized the multi-omics features of 992 long genes (>200 kb), examined their gene length evolution in 15 species, and developed the LongGeneDB to share the multi-omics profiles and the length evolution findings of long genes.

## DATA COLLECTION AND PROCESSING

### Gene annotation data collection and analyses

We analysed the genomic information of 15 species. These species include Homo sapiens (human), Pan troglodytes (chimpanzee), Callithrix jacchus (marmoset), Rattus norvegicus (rat), Mus musculus (mouse), Oryctolagus cuniculus (rabbit), Canis lupus familiaris (dog), Felis catus (cat), Sus scrofa (pig), Taeniopygia guttata (zebra finch), Gallus gallus (chicken), Xenopus tropicalis (frog), Danio rerio (zebrafish), Drosophila melanogaster (fruitfly), and Caenorhabditis elegans (worm). The 15 species belong to 3 Phyla, 6 Classes, 11 Orders, 13 Families, and 15 Genera. The gene annotations of these species were downloaded from the Ensembl Release version 100 (https://useast.ensembl.org/) in gtf file format. In each one of the species, we merged all isoforms of a gene to obtain its entire gene body region and to calculate its gene length. Only protein-coding genes were used for downstream analyses.

### Phylogenetic p-value (phyloP) scores collection

The phyloP scores, which were calculated by the PHAST package for multiple alignments of 59 vertebrate genomes to the mouse genome, were collected from the UCSC Genome Browser (http://hgdownload.cse.ucsc.edu/goldenpath/mm10/phyloP60way/).

### Histone modification ChIP-seq data collection and analyses

The 83 ChIP-seq files of the three histone modifications across the 14 mouse tissues were downloaded from the EBI ENA database (19) (https://www.ebi.ac.uk/ena/browser/home) in sra format using the accession numbers listed in **Supplementary Figure S1**. The ChIP-seq data were generated from 8-week-old mice by the ENCODE project (17). The sra files were converted into FASTQ files by fastq-dump of the SRA Toolkit (https://trace.ncbi.nlm.nih.gov/Traces/sra/sra.cgi?view=software). The FASTQ files were aligned to the mouse mm10 genome by Bowtie version 1.2.3 (20) using the parameters of “-v 2 -m 1 -p 20” to get the sam files. The sam files were converted to bam files by samtools version 1.9 (21). The bam files were further converted to bedgraph files by bamCoverage of deepTools version 3.3.1 (22) using the following parameters: “-of bedgraph -binSize 10 -p 20”.

### Hi-C data collection and analyses

The Hi-C files were downloaded from the EBI ENA database (19) in sra format under the accession numbers of SRX150196 and SRX150197. The Hi-C data were generated from the cerebral cortex of 8-week-old mice by the ENCODE project (17). The sra files were converted into FASTQ files by using fastq-dump of the SRA Toolkit. The FASTQ files were trimmed by using HomerTools (23) (http://homer.ucsd.edu/homer/ngs/homerTools.html) with the following parameters: “trim -3 AAGCTAGCTT -mis 0 -matchStart 20 -min 20”. The trimmed FASTQ files were aligned to the mouse mm10 genome by Bowtie version 1.2.3 (20) to get the sam files. The sam files were used to create the paired-end tag directories by HOMER makeTagDirectory using the following parameters: “-tbp 1 - genome mm10 -restrictionSite AAGCTT -both -removePEbg -removeSelfLigation -removeSpikes 10000 5”. The two biological replicates were merged.

### mRNA-seq data collection and analyses

The 109 mRNA-seq files were downloaded from the EBI ENA database (19) in sra format under the accession numbers listed in **Supplementary Figure S1**. The mRNA-seq data were generated from the 22 tissues of 8-week-old mice by the ENCODE project (17). The sra files were converted into FASTQ files by using fastq-dump of the SRA Toolkit. The FASTQ files were aligned to the mouse mm10 genome by STAR version 2.7.2b (24) with the following parameters: “–runThreadN 13 – outFilterMultimapNmax 1 –outFilterMismatchNmax 3”. The alignment results were saved in sam files. The number of read pairs mapped to the exonic regions of each gene was calculated from the sam files by an in-house Perl script. We calculated the FPKM values by normalizing the raw read counts to the exonic lengths of the gene and to the sequencing depths of the dataset.

### Single nuclear RNA-seq data collection and analyses

The two single nuclear RNA-seq files were downloaded from the EBI ENA database (19) in sra format under the accession numbers of SRR6269025 and SRR6269027. The single nuclear RNA-seq data were generated from 8-week-old mice (25). The sra files were converted into FASTQ files by fastq-dump of the SRA Toolkit. Dropseq-tools V2 (https://github.com/broadinstitute/Drop-seq/releases) was used to process the FASTQ files to get the Digital Gene Expression files. The Digital Gene Expression data were further processed by Seurat 2.3.4 (26) to obtain the expression profiles of the 24 cell types.

## DATABASE CONTENT AND USAGE

### The evolution of gene lengths of long genes in 15 species

Protein-coding genes longer than 200 kb in the mouse genome (992 genes) were included in this database (**Supplementary Figure S1B**). The longest protein isoform of each gene was obtained from Ensembl Release version 100 and was aligned to the protein databases of each of the other 14 species using blastP. The top hit of each alignment was considered to be the ortholog gene. We also used the Ensembl Web Tools of BLAST to align the protein sequences of the 992 mouse genes to the other 14 species to manually confirm the ortholog gene sets. In total, we examined the evolution history of gene lengths of 992 ortholog gene sets in 15 species.

### The omics profiles of long genes in mice

The content of mouse omics profiles for long genes is shown in **Figure 1**. For phyloP scores, we used four subsets of phyloP scores. These scores included glire, euarchontoglire, placental, and all 60 species. Then, we calculated the score profiles of the gene region and the upstream and downstream 100 kb regions of each long gene. To plot the phyloP scores at a single base resolution as bar plots, we used R version 3.6.1. For ChIP-seq, we normalized each profile by their sequencing depth to obtain the reads per million uniquely mapped reads (RPM) values, merged the biological replicates, calculated the merged profiles of the gene region and the upstream and downstream 100 kb regions of each long gene. R was used to plot the ChIP-seq profiles across the 14 tissues as bar plots. For Hi-C, we used analyzeHiC with the following parameters: “-pos” for the gene region and “-res 2000 - window 10000 -balance -cpu 40 -corr”. This calculated the Pearson’s correlation coefficient of normalized Hi-C interactions for the gene region and the upstream and downstream 100 kb regions of each long gene. Then, we used R to plot the matrix as a heatmap. For mRNA-seq, we obtained the FPKM values across the 22 tissues for each long gene and used R to plot them as a boxplot. For single nuclear RNA-seq, we used R to plot the expression levels of long genes and three cell-type-marker genes in the 24 cell types in violin plots.

**Figure 1.**
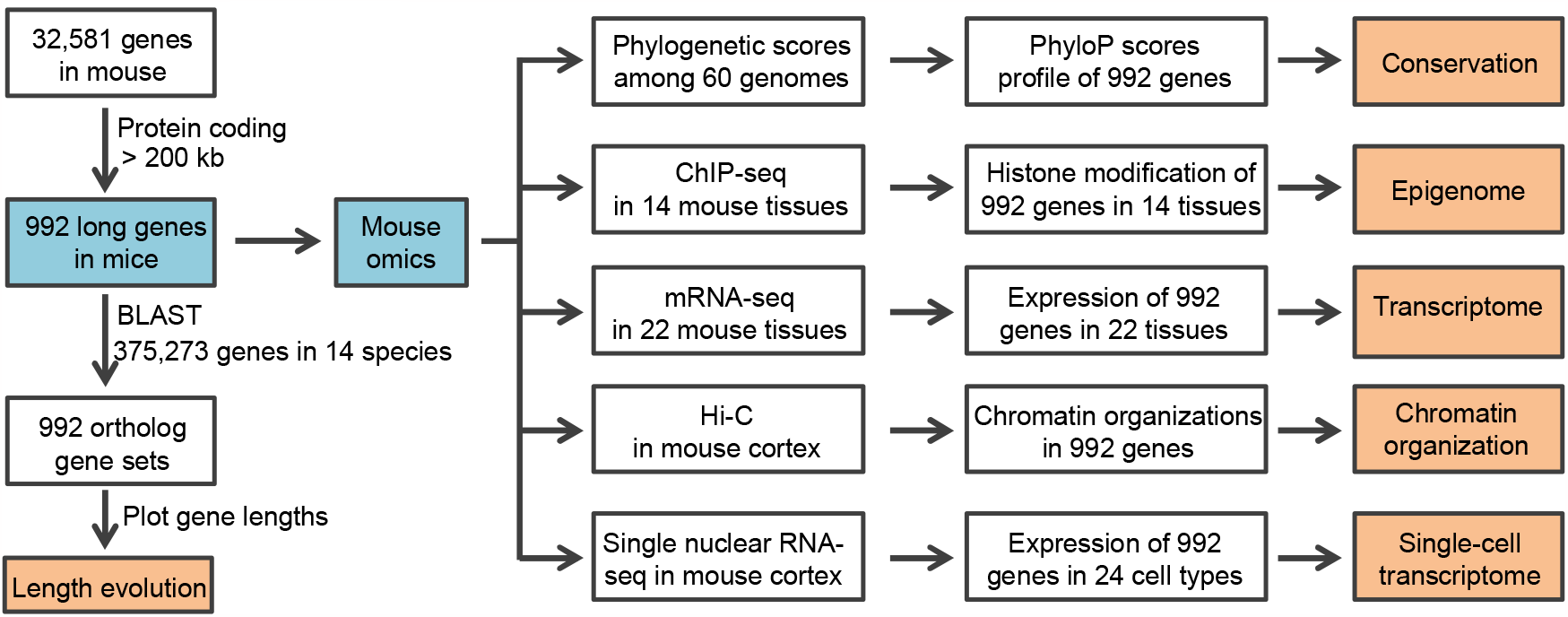
Construction pipeline of the LongGeneDB database. Gene annotation files were collected from the Ensembl database. Protein-coding genes that are longer than 200 kb in the mouse genome were selected. The blastP was used to identify their orthologous genes in the other 14 species. The phyloP scores were collected from the UCSC Genome Browser. The raw data of ChIP-seq, mRNA-seq, Hi-C, and single nuclear RNA-seq were collected from the EBI ENA database. Users can query the pages of Length evolution and Mouse omics and download all the plots and their source data.

### Web design and interface

LongGeneDB provides an easily accessible web interface developed in the web server stack of the Linux operating system, the Apache HTTP Server, the MySQL database management system, and the PHP programming language (LAMP). The dynamic website was created by PHP, HTML, CSS, and JavaScript code and other scripts. Plotting data was performed by using R version 3.6.1. The LongGeneDB contains two components, the Length Evolution and the Mouse Omics (**Figure 2A**). In each component, users can easily query LongGeneDB by typing the gene symbol in the search box (**Figure 2B**). The search box is case-insensitive.

**Figure 2.**
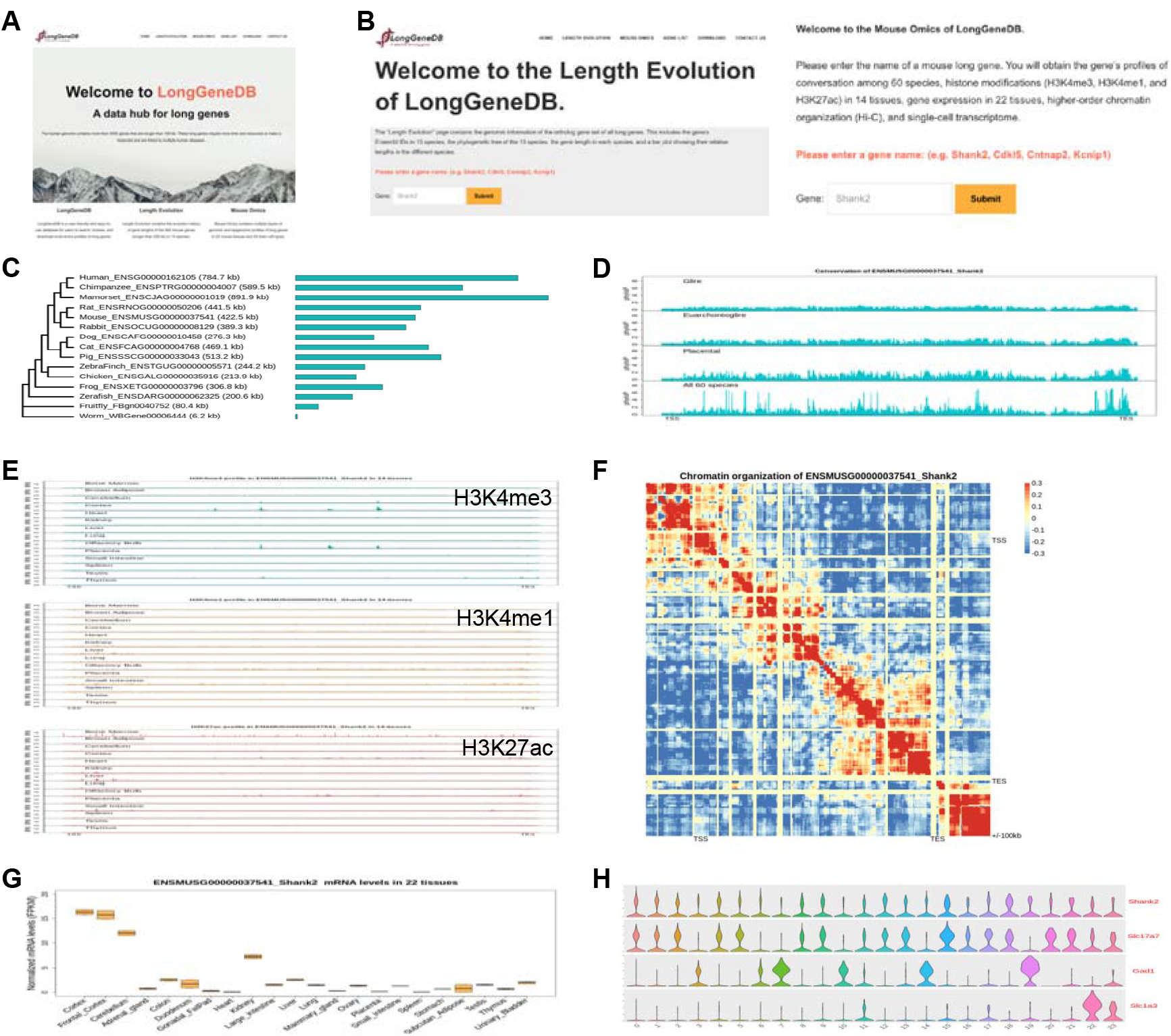
Overview of LongGeneDB. (**A**) The home page of the LongGeneDB. (**B**) Query interface of the Length Evolution page (left) and the Mouse Omics page (right). Users can search the two components by entering a gene symbol. (**C**) A bar plot shows the evolution of the gene length of Shank2 in 15 species. (**D**) Bar plots show the conservation of the mouse Shank2 gene region. (**E**) Bar plots show the profiles of the three types of histone modifications in Shank2 gene region in the 14 mouse tissues. (**F**) A heatmap shows the chromatin organization in the gene body region of Shank2. (**G**) A boxplot shows the expression levels of Shank2 in the 22 mouse tissues. (**H**) Violin plots show the expression levels of Shank2 and three cell-type-marker genes in the 24 cell types in the mouse cerebral cortex.

### Data browsing and querying

Query on the “Length Evolution” page returns the genomic information of the ortholog gene set of the queried gene. This includes the gene Ensembl IDs in 15 species, the phylogenetic tree of the 15 species, the gene length in each species, and a bar plot showing their relative lengths in the different species. For example, querying the gene “*Shank2*” returned the evolutionary history of the gene lengths of the *Shank2* ortholog gene set (**Figure 2C**). The gene region of *Shank2* is shown to become longer during evolution.

The query on the “Mouse Omics” page returns mouse multi-omics profiles of the queried gene. The results include bar plots of its conservation in gene bodies across 60 vertebrate genomes, bar plots of the profiles of the three types of histone modifications in the 14 tissues, a heatmap of the chromatin organization in its gene body region, a boxplot of its expression profile in the 22 tissues, and violin plots of its expression profile in the 24 cell types in the mouse cortex. For example, querying the gene “*Shank2*” showed that the *Shank2* gene region contains 11 conserved regions (**Figure 2D**). *Shank2* is actively transcribed in three regions of the brain and has four alternative promoters (**Figure 2E**). The chromatin in the gene body of *Shank2* is organized into two domains (**Figure 2F**). *Shank2* is highly expressed in three regions of the brain (**Figure 2G**) and is constitutively expressed in the 24 cell types in the mouse cerebral cortex (**Figure 2H**).

The “Gene List” page provides the mouse gene symbols of the 992 long genes. The “Help” page provides the scientific classifications of the 15 species, the analysis pipelines, and the accession numbers for the omics data.

### Download data and figures

All figures generated in LongGeneDB can be downloaded in PNG format by right-clicking the figure of interest and choosing “Save Image As…”. On the ‘Download’ page, users can download the source data files in gzip format. The source data includes the gene lengths of all genes in the 15 species, all the scripts used in this study, the lengths of the 992 ortholog gene sets in the species, the conservation scores, the ChIP-seq values of H3K4me3, H3K4me1, and H3K27ac, the gene expression FPKM values in the 22 tissues, the Hi-C contact matrix, and the single nuclear RNA-seq data. Users can also download all the plots of the 992 ortholog gene sets in PNG format in tar.gz files.

## SUMMARY

We developed the LongGeneDB as an easy-to-use database for querying and downloading multiomics profiles of long genes. By integrating large-scale omics datasets, LongGeneDB provides a comprehensive multi-omics profiles of 992 ortholog long gene sets in 15 species and across 22 mouse tissues. LongGeneDB includes protein-coding genes that are longer than 200 kb in the mouse genome, and we will expand the gene list by including genes that are longer than 100 kb in one of the 15 species. We will update the LongGeneDB to include disease-related multi-omics data from studies in humans, such as data from the Genomic Data Commons Data Portal (27) and the PsychENCODE project (28). Since the number of human diseases that are associated with long genes is constantly increasing, it is necessary to further illuminate the molecular mechanisms underlying long genes. We will maintain LongGeneDB as a useful resource for the biomedical research community to understand long genes.

## SUPPLEMENTARY DATA

Supplementary Data are available at ### online.

## REFERENCES

1. Yates, A.D., Achuthan, P., Akanni, W., Allen, J., Allen, J., Alvarez-Jarreta, J., Amode, M.R., Armean, I.M., Azov, A.G., Bennett, R. et al. (2020) Ensembl 2020. Nucleic Acids Res, 48, D682–D688.

2. Polymenidou, M., Lagier-Tourenne, C., Hutt, K.R., Huelga, S.C., Moran, J., Liang, T.Y., Ling, S.C., Sun, E., Wancewicz, E., Mazur, C. et al. (2011) Long pre-mRNA depletion and RNA missplicing contribute to neuronal vulnerability from loss of TDP-43. Nat Neurosci, 14, 459–468.

3. Lagier-Tourenne, C., Polymenidou, M., Hutt, K.R., Vu, A.Q., Baughn, M., Huelga, S.C., Clutario, K.M., Ling, S.C., Liang, T.Y., Mazur, C. et al. (2012) Divergent roles of ALS-linked proteins FUS/TLS and TDP-43 intersect in processing long pre-mRNAs. Nat Neurosci, 15, 1488–1497.

4. King, I.F., Yandava, C.N., Mabb, A.M., Hsiao, J.S., Huang, H.S., Pearson, B.L., Calabrese, J.M., Starmer, J., Parker, J.S., Magnuson, T. et al. (2013) Topoisomerases facilitate transcription of long genes linked to autism. Nature, 501, 58–62.

5. Sugino, K., Hempel, C.M., Okaty, B.W., Arnson, H.A., Kato, S., Dani, V.S. and Nelson, S.B. (2014) Cell-type-specific repression by methyl-CpG-binding protein 2 is biased toward long genes. J Neurosci, 34, 12877–12883.

6. Gabel, H.W., Kinde, B., Stroud, H., Gilbert, C.S., Harmin, D.A., Kastan, N.R., Hemberg, M., Ebert, D.H. and Greenberg, M.E. (2015) Disruption of DNA-methylation-dependent long gene repression in Rett syndrome. Nature, 522, 89–93.

7. Johnson, B.S., Zhao, Y.T., Fasolino, M., Lamonica, J.M., Kim, Y.J., Georgakilas, G., Wood, K.H., Bu, D., Cui, Y., Goffin, D. et al. (2017) Biotin tagging of MeCP2 in mice reveals contextual insights into the Rett syndrome transcriptome. Nat Med, 23, 1203–1214.

8. Takeuchi, A., Iida, K., Tsubota, T., Hosokawa, M., Denawa, M., Brown, J.B., Ninomiya, K., Ito, M., Kimura, H., Abe, T. et al. (2018) Loss of Sfpq Causes Long-Gene Transcriptopathy in the Brain. Cell Rep, 23, 1326–1341.

9. Zhao, Y.T., Kwon, D.Y., Johnson, B.S., Fasolino, M., Lamonica, J.M., Kim, Y.J., Zhao, B.S., He, C., Vahedi, G., Kim, T.H. et al. (2018) Long genes linked to autism spectrum disorders harbor broad enhancer-like chromatin domains. Genome Res, 28, 933–942.

10. Barbash, S. and Sakmar, T.P. (2017) Length-dependent gene misexpression is associated with Alzheimer’s disease progression. Sci Rep, 7, 190.

11. Helmrich, A., Stout-Weider, K., Hermann, K., Schrock, E. and Heiden, T. (2006) Common fragile sites are conserved features of human and mouse chromosomes and relate to large active genes. Genome Res, 16, 1222–1230.

12. Smith, D.I., McAvoy, S., Zhu, Y. and Perez, D.S. (2007) Large common fragile site genes and cancer. Semin Cancer Biol, 17, 31–41.

13. Helmrich, A., Ballarino, M. and Tora, L. (2011) Collisions between replication and transcription complexes cause common fragile site instability at the longest human genes. Mol Cell, 44, 966–977.

14. Manzo, S.G., Hartono, S.R., Sanz, L.A., Marinello, J., De Biasi, S., Cossarizza, A., Capranico, G. and Chedin, F. (2018) DNA Topoisomerase I differentially modulates R-loops across the human genome. Genome Biol, 19, 100.

15. Wei, P.C., Chang, A.N., Kao, J., Du, Z., Meyers, R.M., Alt, F.W. and Schwer, B. (2016) Long Neural Genes Harbor Recurrent DNA Break Clusters in Neural Stem/Progenitor Cells. Cell, 164, 644–655.

16. Wang, R., Zheng, D., Wei, L., Ding, Q. and Tian, B. (2019) Regulation of Intronic Polyadenylation by PCF11 Impacts mRNA Expression of Long Genes. Cell Rep, 26, 2766–2778 e2766.

17. Consortium, E.P., Moore, J.E., Purcaro, M.J., Pratt, H.E., Epstein, C.B., Shoresh, N., Adrian, J., Kawli, T., Davis, C.A., Dobin, A. et al. (2020) Expanded encyclopaedias of DNA elements in the human and mouse genomes. Nature, 583, 699–710.

18. Barrett, T., Wilhite, S.E., Ledoux, P., Evangelista, C., Kim, I.F., Tomashevsky, M., Marshall, K.A., Phillippy, K.H., Sherman, P.M., Holko, M. et al. (2013) NCBI GEO: archive for functional genomics data sets--update. Nucleic Acids Res, 41, D991–995.

19. Amid, C., Alako, B.T.F., Balavenkataraman Kadhirvelu, V., Burdett, T., Burgin, J., Fan, J., Harrison, P.W., Holt, S., Hussein, A., Ivanov, E. et al. (2020) The European Nucleotide Archive in 2019. Nucleic Acids Res, 48, D70–D76.

20. Langmead, B., Trapnell, C., Pop, M. and Salzberg, S.L. (2009) Ultrafast and memory-efficient alignment of short DNA sequences to the human genome. Genome Biol, 10, R25.

21. Li, H., Handsaker, B., Wysoker, A., Fennell, T., Ruan, J., Homer, N., Marth, G., Abecasis, G., Durbin, R. and Genome Project Data Processing, S. (2009) The Sequence Alignment/Map format and SAMtools. Bioinformatics, 25, 2078–2079.

22. Ramirez, F., Dundar, F., Diehl, S., Gruning, B.A. and Manke, T. (2014) deepTools: a flexible platform for exploring deep-sequencing data. Nucleic Acids Res, 42, W187–191.

23. Heinz, S., Benner, C., Spann, N., Bertolino, E., Lin, Y.C., Laslo, P., Cheng, J.X., Murre, C., Singh, H. and Glass, C.K. (2010) Simple combinations of lineage-determining transcription factors prime cis-regulatory elements required for macrophage and B cell identities. Mol Cell, 38, 576–589.

24. Dobin, A., Davis, C.A., Schlesinger, F., Drenkow, J., Zaleski, C., Jha, S., Batut, P., Chaisson, M. and Gingeras, T.R. (2013) STAR: ultrafast universal RNA-seq aligner. Bioinformatics, 29, 15–21.

25. Hu, P., Fabyanic, E., Kwon, D.Y., Tang, S., Zhou, Z. and Wu, H. (2017) Dissecting Cell-Type Composition and Activity-Dependent Transcriptional State in Mammalian Brains by Massively Parallel Single-Nucleus RNA-Seq. Mol Cell, 68, 1006–1015 e1007.

26. Butler, A., Hoffman, P., Smibert, P., Papalexi, E. and Satija, R. (2018) Integrating single-cell transcriptomic data across different conditions, technologies, and species. Nat Biotechnol, 36, 411–420.

27. Grossman, R.L., Heath, A.P., Ferretti, V., Varmus, H.E., Lowy, D.R., Kibbe, W.A. and Staudt, L.M. (2016) Toward a Shared Vision for Cancer Genomic Data. N Engl J Med, 375, 1109–1112.

28. Psych, E.C., Akbarian, S., Liu, C., Knowles, J.A., Vaccarino, F.M., Farnham, P.J., Crawford, G.E., Jaffe, A.E., Pinto, D., Dracheva, S. et al. (2015) The PsychENCODE project. Nat Neurosci, 18, 1707–1712.

